# Explainable Multimodal Machine Learning Using Combined Environmental DNA and Biogeographic Features for Ecosystem Biomonitoring

**DOI:** 10.1101/2025.05.07.652781

**Authors:** Joshua C.O. Koh

**Affiliations:** Office of the Supervising Scientist, Eaton, Northern Territory 0820, Australia

**Keywords:** Environmental DNA, amplicon sequence variants, operational taxonomic unit, automated machine learning, differential abundance analysis, ecosystem biomonitoring

## Abstract

Machine learning (ML) has been proposed as a taxonomy-independent approach using environmental DNA (eDNA) for ecosystem biomonitoring. Representations of eDNA amplicons either as clustered sequences termed operational taxonomic units (OTUs) or unique sequences termed amplicon sequence variants (ASVs) are used as inputs in current ML practices, with varied successes reported. Biogeographic data encompassing the physical, climate and ecological observations collected at biomonitoring sites provide a repository of potential informative features that can augment ML performance in combination with eDNA data. I introduce a multimodal ML workflow using combined eDNA and biogeographic features for ecosystem biomonitoring. Differentially abundant ASVs were merged with biogeographic data and used as input in an automated ML approach. Using Switzerland’s freshwater macroinvertebrate eDNA dataset collected across 163 biomonitoring sites and impact prediction as an example, I show that the multimodal ML approach (83.3% accuracy) significantly outperformed ML using only ASVs (66.7% accuracy) or OTUs (64.6% accuracy). Shapley additive explanation of the best ML model revealed key biogeographic features and species/taxa impacting upon predictions. The proposed workflow can be readily adopted in existing bioinformatics/ML pipelines and will further advance the use of eDNA for biomonitoring.

## INTRODUCTION

Recent global reports ^1, 2^ have painted a bleak outlook for biodiversity with anthropogenic activities and accelerating climate change driving an unprecedented decline in biodiversity. Experts estimate about 30% of species have been globally threatened or driven to extinction since the year 1500, with this number increasing to 37% by 2100 if the trend persists ^3^. Central to the success of environmental management and conservation efforts are effective ecosystem monitoring strategies that can detect or predict early critical shifts in the environment ^3^. One of the emerging monitoring tools is DNA-based biomonitoring, specifically environmental DNA (eDNA) metabarcoding which seeks to identify multiple taxa concurrently from environmental samples using standard genetic markers ^4, 5^. Compared to manual approaches, eDNA metabarcoding offers a non-invasive, high-throughput and cost-effective method to retrieve biodiversity inventories across a broad spatiotemporal scale ^4, 5^. Nonetheless, the success of eDNA metabarcoding is dependent upon the availability of reference sequences for taxonomic annotation and the method suffers from biases in PCR (polymerase chain reaction) amplification and sequencing ^4, 6^. In addition, studies comparing eDNA-based and traditional morphology-based taxonomic inventories have shown varied degrees of congruence ^7^.

In contrast to the taxonomy-based approach, the de novo approach seeks to directly associate biological community profiles (including identified and unidentified taxa) retrieved via eDNA metabarcoding with known ecological status or disturbance gradients ^4^. This approach promises to harness the full potential of high-throughput genetic data and move towards a more holistic monitoring paradigm ^4, 8^. To this end, machine learning (ML) has emerged as a promising taxonomy-independent approach for ecosystem biomonitoring using eDNA, with its ability to capture complex non-linear relationships between multiple environmental pressures and the biological diversity provided by metabarcoding ^9, 10^. Early studies focused on using ML to predict biological indices traditionally obtained using manual approaches in both marine ^11, 12^ and freshwater ^9, 13^ ecosystems, with examples of ML-inferred index values outperforming the taxonomy-based strategy ^11, 12^. However, the use of ML to infer traditional indices also inherits the errors and biases present in these indices and limits the added value of eDNA for ecological assessment ^8^. A recent study has attempted to circumvent these constraints by directly inferring environmental states using features present in eDNA with ML ^8^. Using impact (due to agriculture and urbanization) prediction for 64 federal monitoring sites in Switzerland as an example, Keck *et al.*^8^ showed that a random forest model trained on freshwater macroinvertebrate eDNA performed better (69.1% accuracy) compared to a model trained on morpho-taxonomy data obtained via traditional kicknet sampling (59.5% accuracy), suggesting that information contained in eDNA alone can be used to infer environmental states at the landscape scale.

Nevertheless, current ML practices that use eDNA or genetic data as the sole input in model training inherently limit the true potential of the ML approach for ecosystem biomonitoring as it is unlikely that metabarcoding alone can fully account for the complex drivers responsible for environmental change ^14, 15^. Ecosystem modelling, a discipline wherein quantitative and qualitative models are used to disentangle interactions between ecosystem components and make predictions about future ecosystem states, commonly employ multiple data modalities/types (e.g., geomorphological, climate, hydrology, biological and genetic/genomic data) in the model building process ^15^. In contrast, such practice has yet to be adopted in ML approaches using eDNA for ecosystem biomonitoring. In that regard, biogeographic data (e.g., geological, ecological, climate and biological factors that impact on the spatiotemporal distribution of organisms) collected at biomonitoring sites serves as an untapped repository of potential high-quality features that can augment ML performance in combination with eDNA/genetic data. However, the high dimensionality (where features or sequence data greatly outnumber samples) of eDNA metabarcoding data poses a technical challenge for a multimodal ML implementation as signals from introduced features (e.g., biogeographic data) may be lost as noise or obscured in the enormity of sequence data. In addition, a multimodal dataset with mixed data types, for example, quantitative (discrete and continuous numeric data) and qualitative (categorical text and ordinal text/numeric data) features, require careful data representation to ensure that different modalities can be integrated in a meaningful manner. It is likely that an exploration of ML models beyond the customary random forest and/or support vector machine ^12, 16^ is required to uncover the best model architecture(s) amenable to multimodal datasets.

In this study, I introduce a multimodal ML workflow using combined eDNA and biogeographic data for ecosystem biomonitoring. The workflow consists of modular components that collectively enhance the reproducibility, performance and explainability of the ML approach. Using Switzerland’s freshwater macroinvertebrate eDNA dataset collected across 163 federal biomonitoring sites and impact prediction as an example, I demonstrate the application of the multimodal ML workflow using six popular ML models and compare its performance against ML using only eDNA. The proposed workflow is compatible with existing bioinformatics/ML pipelines and provides insights into species/taxa and biophysical factors impacting upon environmental states using model explanation techniques in conjunction with taxonomic information.

## MATERIAL AND METHODS

### eDNA Metabarcoding Data and Site Classification

Data used in this study were derived from 163 eDNA sampling sites (Figure 1) which were part of a Swiss national surface water quality biomonitoring program (NAWA) ^17^ and available publicly ^8^. These sites had metadata for biogeographic information, specifically the corresponding biogeographic region ^18^, catchment area and site altitude. Sampling occurred in the summer of 2018 and the spring of 2019, with a total of four eDNA water samples collected per site and DNA was extracted as described elsewhere ^8^. The DNA samples were amplified using the fwhF/EPTDr2n primer pair which targets a 142 bp fragment of the COI (cytochrome C oxidase subunit I) marker ^19, 20^. A COI sequencing library was prepared and sequenced (paired-end) on an Illumina MiSeq ^8^. Sites were classified as reference if their proportion of natural land use (i.e., surfaces not classified as agriculture or urban areas, mainly forests) was higher than 33% ^8^ and as impacted otherwise, resulting in 83 reference and 80 impacted sites (Figure 1).

**Figure 1.**
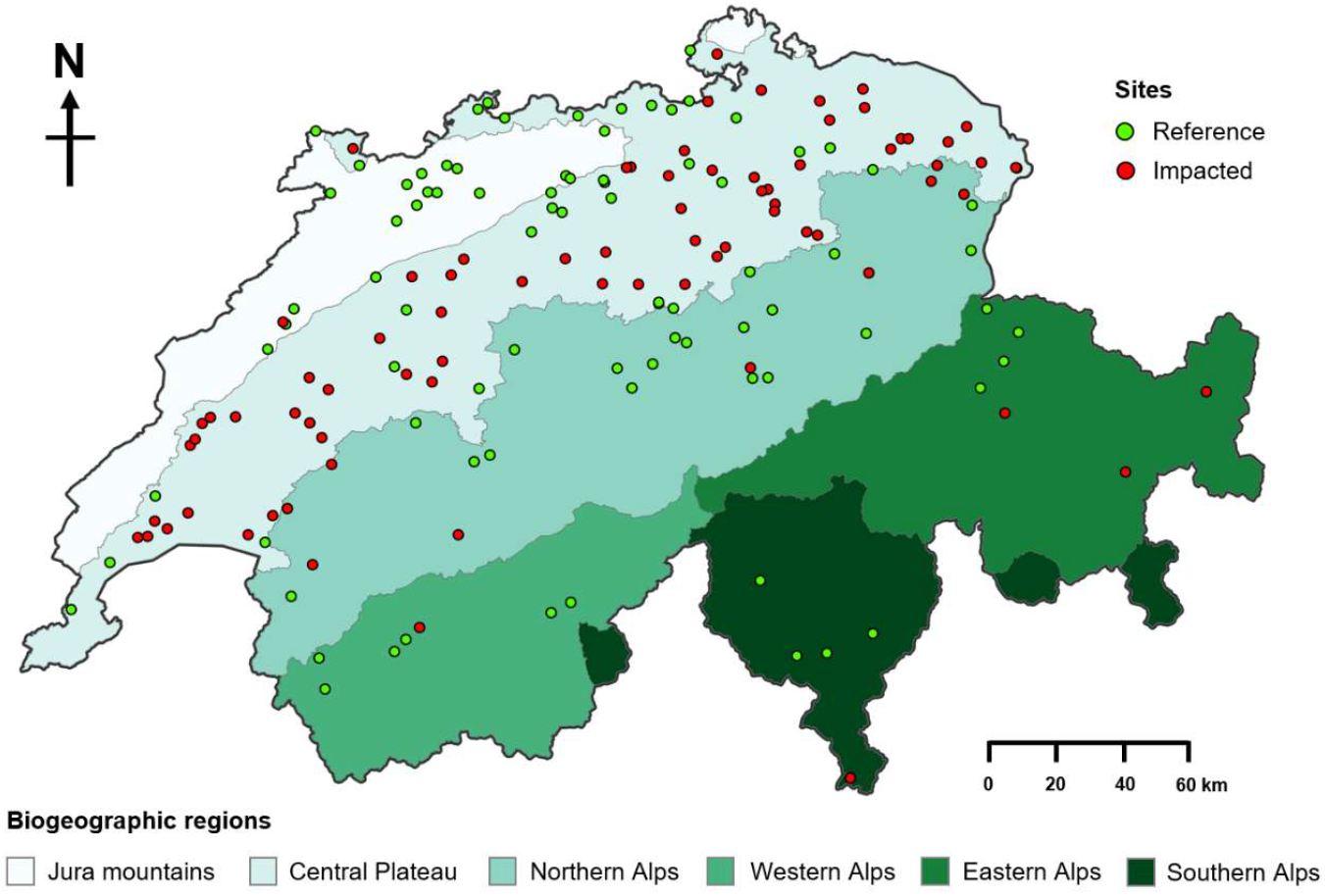
Map of Switzerland showing the location of the 163 biomonitoring sites distributed across six biogeographic regions.

### Bioinformatics

Processing of the COI library to yield non-chimeric amplified sequence variants (ASVs) based on the DADA2 R package ^21^ and subsequent taxonomic annotation using a custom macroinvertebrate database are described in Keck *et al.* ^8^. Both the ASV read count table, and the taxonomic annotations were used as inputs in this study. Rare ASVs (<10 occurrence) were removed and only those with sequence length more than 137 bp and shorter than 147 bp were retained. A final filtering step was performed to remove ASVs that were found in relative proportion more than 0.1% in one of the negative controls (i.e., negative extraction). The resulting ASVs were clustered into OTUs (operational taxonomic units) using VSEARCH version 2.27 with default settings ^22^. Each OTU is represented by a consensus sequence, which is the most common or representative sequence within a cluster of sequences based on a similarity threshold (97% sequence identity). Differential abundance (DA) analysis was performed on the ASVs using the ANCOM-BC2 R package version 2.8.1 ^23^ and the standalone differential ranking (DR) package, Songbird, version 1.0.4 ^24^, with the site status (reference or impacted) specified as the condition for the statistical testing. For ANCOM-BC2, default settings were used; for DR, the following parameters were used: epochs=5000, differential-prior=0.5, min-feature-count=10 and min-sample-count=100. Taxonomic annotations for ASVs identified as important features for impact prediction (see below) were verified by subjecting the sequences to BLASTN (Basic Local Alignment Search Tool Nucleotide) search against NCBI (National Centre for Biotechnology Information) online nucleotide database, with previous annotations ^8^ that had low assignment confidence (e.g., <30%) replaced with those from BLAST hits that had high sequence identity and coverage (e.g., > 90%) (Supporting Information Table S1).

### Multimodal Machine Learning Workflow

A multimodal ML workflow consisting of five steps: differential abundance analysis, feature filtering, feature merging, automated ML and model explanation was developed to facilitate ML using combined eDNA and biogeographic data to directly infer environmental states (Figure 2). The workflow adopts ASVs as the standard unit of eDNA analysis (although it is also compatible with other eDNA representations, e.g., OTU) to enhance reusability and reproducibility of the method, as ASVs being unique sequences (down to single-nucleotide difference) provide absolute and consistent labels transferable across different experiments ^25^. In addition, ASVs offer the finest taxonomic resolution (down to species level) where reference sequences are available, and a recent study showed that ML models trained at the highest taxonomic resolution (ASVs) were most accurate in predicting measures of soil health ^26^.

**Figure 2.**
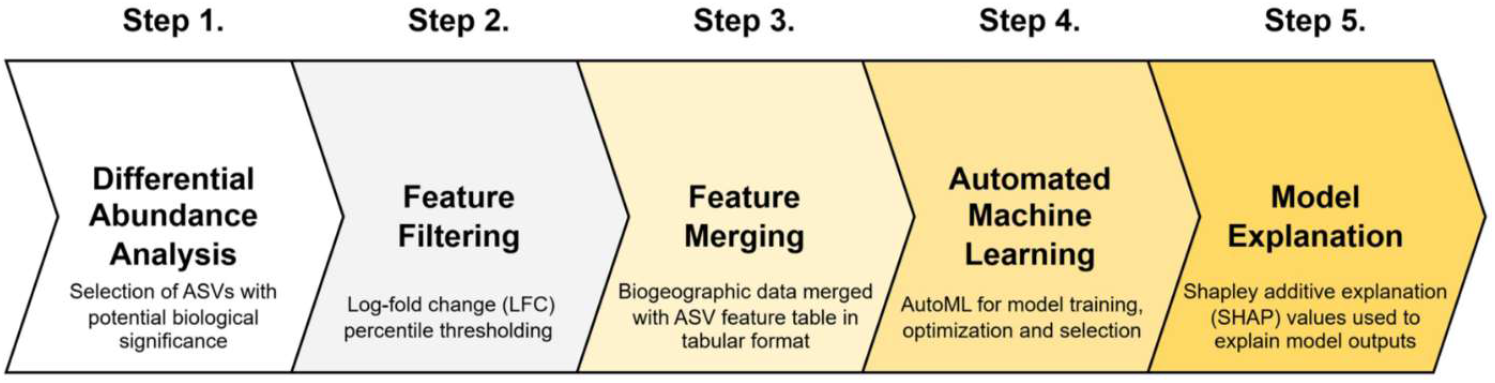
Multimodal machine learning workflow.

However, the enormity of ASV sequence data, as opposed to clustered sequences such as OTUs (i.e., reduced dimensionality) presents computational challenges/bottlenecks and is not readily amenable to combining with other data sources/types. Although feature selection methods common in ML practices (e.g., uni/multivariate statistics-based filtering, recursive forward/backward feature selection, and feature importance embedded in models) have been proposed to address issues concerned with the high dimensionality of ASV data ^27^, these generic methods do not consider the compositional nature of eDNA metabarcoding data and are not designed to identify features responding specifically to biological/ecological pressures, whilst controlling for false detections and sequencing biases.

In contrast, recent DA analyses based on compositional statistics were effective in identifying biologically significant features/taxa from microbiomes with examples in environmental and human medical case studies with low false discoveries ^23, 24, 28^. Thus, DA analysis was selected as the dimension reduction/feature selection strategy (Step 1, Figure 2) in the workflow. Two popular DA analysis methods, ANCOM-BC2 ^23^ and differential ranking (DR) ^24^, were implemented in this study for ASV selection. The selected ASVs were further filtered (Step 2, Figure 2) using a percentile log-fold change (LFC, see section below) thresholding to retain ASVs with high differentials (e.g., those potentially with the strongest signals). The resulting list (or membership) of ASVs serves as an absolute reference for the construction of a feature table suited for feature merging (Step 3, Figure 2) where selected ASVs’ (columns) read counts are summed according to samples (rows) and normalized within sample to the range of 0 to 1 using the MinMaxScaler function in the Scikit-learn Python package version 1.4 ^29^. Biogeographic data (text or numeric) were then appended directly without modification as additional columns to the resulting ASV feature table.

The combined feature table was used as input for ML model training via an automated machine learning (AutoML) approach (Step 4, Figure 2) which streamlines the entire process of data preparation (cleaning and preprocessing), feature engineering, model generation (selection and optimization), and model evaluation with minimal effort/intervention from the user ^30^. The use of AutoML systems not only allows rapid prototyping of multiple ML models for classification (e.g., assigning labels) or regression (e.g., predicting values) tasks but also automates the complex task of feature encoding to account for the different data types present in multimodal datasets. In this study, the PyCaret Python package version 3.0 ^31^ was implemented as the AutoML system of choice as it provides a balanced approach to ML automation (compared to fully turnkey systems), providing the user with the option to partially or fully define components within the AutoML pipeline. In addition, PyCaret outputs complete ML pipelines in Scikit-learn format, which is widely considered the ML standard in Python, allowing for easy sharing and ensuring reproducibility and transparency of the ML workflows. Six popular ML models ^30, 32, 33^: support vector machine with radial basis function kernel (SVM), random forests (RF), light gradient boosted machine (LightGBM), extreme gradient boosting (XGBoost), categorical boosting (CatBoost) and multilayer perceptron (MLP, a type of artificial neural network) were implemented using PyCaret. Of these, RF, LightGBM, XGBoost and CatBoost are collectively referenced as tree-based models as they use decision trees to make predictions ^33^.

The final step in the workflow is modal explanation/interpretability (Step 5, Figure 2), which seeks to trace the decision-making process of a ML model and understand the key features impacting upon its predictions ^34^. This is crucial to inform decision making in environmental regulatory/management settings and instill trust in the ML tools. Shapley additive explanations (SHAP), a popular feature-based model explanation method ^35^ was implemented in the workflow using the SHAP Python package version 0.46. The best ML model(s) discovered in AutoML was subjected to SHAP analysis, which computes individual contributions of each feature (i.e., SHAP values) to the predictions based on game theory ^35^. SHAP values provide insight into a model’s behavior either across the entire dataset (global explanation, e.g., this study) or for specific predictions (local explanation). In this study, SHAP values across the training set were derived using the model-agnostic KernelExplainer method, with SHAP values for the top ten features visualized in a bar plot indicating feature importance and in a beeswarm plot showing the effect of individual features on predictions, i.e., how changes in feature values affect predictions ^36^.

### Multimodal Machine Learning for Impact Prediction

A binary classification task predicting site status as either reference (Ref) or impacted (Imp) for 163 eDNA biomonitoring sites described previously was used to demonstrate and evaluate the performance of the proposed multimodal ML workflow. The eDNA samples were collected from sites distributed across six biogeographic regions and from two seasons (89 sites had samples from one season; 74 sites had samples from two seasons), providing a diverse and heterogeneous dataset for testing. To account for potential seasonal effects, sites were further coded as site-season (e.g., SPEZ_033_FR-spring, SPEZ_033_FR-summer) for ML, resulting in a total of 237 site-season samples which were split randomly in a ratio of 80:20 into training (n=189) and test (n=48) sets, with both datasets having the same site status distribution (i.e., stratified sampling). The resulting training and test sets (i.e., the same site-season splits) were used for all ML experiments to ensure comparability. The ML experiments were set up in PyCaret using the ClassificationExperiment class, with the six ML models specified previously trained on the training set using a 10-fold cross-validation strategy ^37^ and with model hyperparameter optimization enabled using the Optuna tuner ^38^ with 100 iterations or optimization rounds. Model performance was evaluated on the test set (i.e., unseen data) using classification accuracy (proportion of correctly classified samples out of all samples), with additional metrics such as precision (false positive indicator; high precision=low false positives), recall (false negative indicator; high recall=low false negatives) and F1-score (combined metric of precision and recall) provided for the best model from each ML model (Supporting Information Table S2).

The effect of different dimension reduction and/or feature selection methods (e.g., OTU clustering and DA analysis) for ASVs on ML model performance was first determined by comparing classification accuracy of the six ML models across four datasets: ASVs (i.e., total ASVs), OTUs, ASVs selected by ANCOM-BC2 (ASV_ANCOM-BC2) and ASVs selected by DR (ASV_DR). The feature table for each dataset was constructed by summing total read counts of each feature (ASV or OTU in columns) according to site-season samples (rows) and normalized to the range of 0 to 1 as described previously within samples. Normalization was performed within each sample as opposed to between samples to ensure that the operation is dataset-independent (i.e., not dependent on the feature values across a specific dataset) and transferable across datasets. Next, the effect of further filtering of differentially abundant ASVs based on the log-fold change (LFC) on ML performance was examined using the DA method which yielded the best ML performance. The LFC is a metric in DA analysis which quantifies the magnitude and direction (negative or positive values) of change in the abundance of a feature (e.g., ASV) between two conditions (e.g., Ref and Imp) on a logarithmic scale ^28^. Differentially abundant ASVs were filtered using the 30^th^, 40^th^ and 50^th^ percentile of the absolute LFC values as minimum thresholds, and the performance of the top two ML models identified from the previous ML experiment was evaluated using the filtered ASV datasets. The ML experiment was then repeated for all six ML models using the optimum filtered ASV dataset, and SHAP analysis was performed on the best ML model with taxonomic annotations displayed where available (Supporting Information Table S1) in visualizations.

Lastly, a combined multimodal dataset for AutoML was generated as described previously by merging the filtered ASV feature table with biogeographic data where information for altitude, biogeographic region and catchment was appended as additional columns. SHAP analysis was performed on all ML models to understand the impact of the added biogeographic data on model performance. Both SHAP bar and beeswarm plots were generated for the best ML model.

## RESULTS

### Impact prediction using eDNA data

The effect of ASVs dimension reduction (OTU clustering) and feature selection (DA analysis) on ML performance was first determined (Figure 3A). Clustering of 23,628 ASVs yielded 6,465 OTUs, representing a 72.6% reduction in data size. In comparison, DA analyses produced the most drastic reduction in data size, with ANCOM-BC2 identifying 293 ASVs (98.8 % reduction) as differentially abundant across reference and impacted sites, followed by DR with 1,339 ASVs (94.3 % reduction) identified as differentially abundant. Interestingly, all 293 ASVs identified by ANCOM-BC2 were present in the 1,339 ASVs identified by DR, suggesting that both methods were congruent with ANCOM-BC2 imposing a more stringent selection on ASVs. Overall ML classification performance (% accuracy) was comparable across four datasets: ASV (50.00 – 66.67%), OTU (50.00 – 64.58%), ASV_ANCOM-BC2 (54.17 – 64.58%) and ASV_DR (56.25 – 68.75%), with the best performance (68.75% accuracy) produced by a MLP model trained on the ASV_DR dataset (Figure 3A). These results indicate that both OTU clustering and DA analysis can be effective dimension reduction methods without adversely impacting on ML performance, albeit DA analysis (DR) offered enhanced ML performance in addition to selecting ASVs with potential biological/ecological significance in this study. Between the six ML models, a clear trend can be observed across all four datasets, wherein the best ML performance was registered by the MLP model, followed by CatBoost as second best, and the rest generally in the order of XGBoost, LightGBM, RF and SVM (Figure 3A).

**Figure 3.**
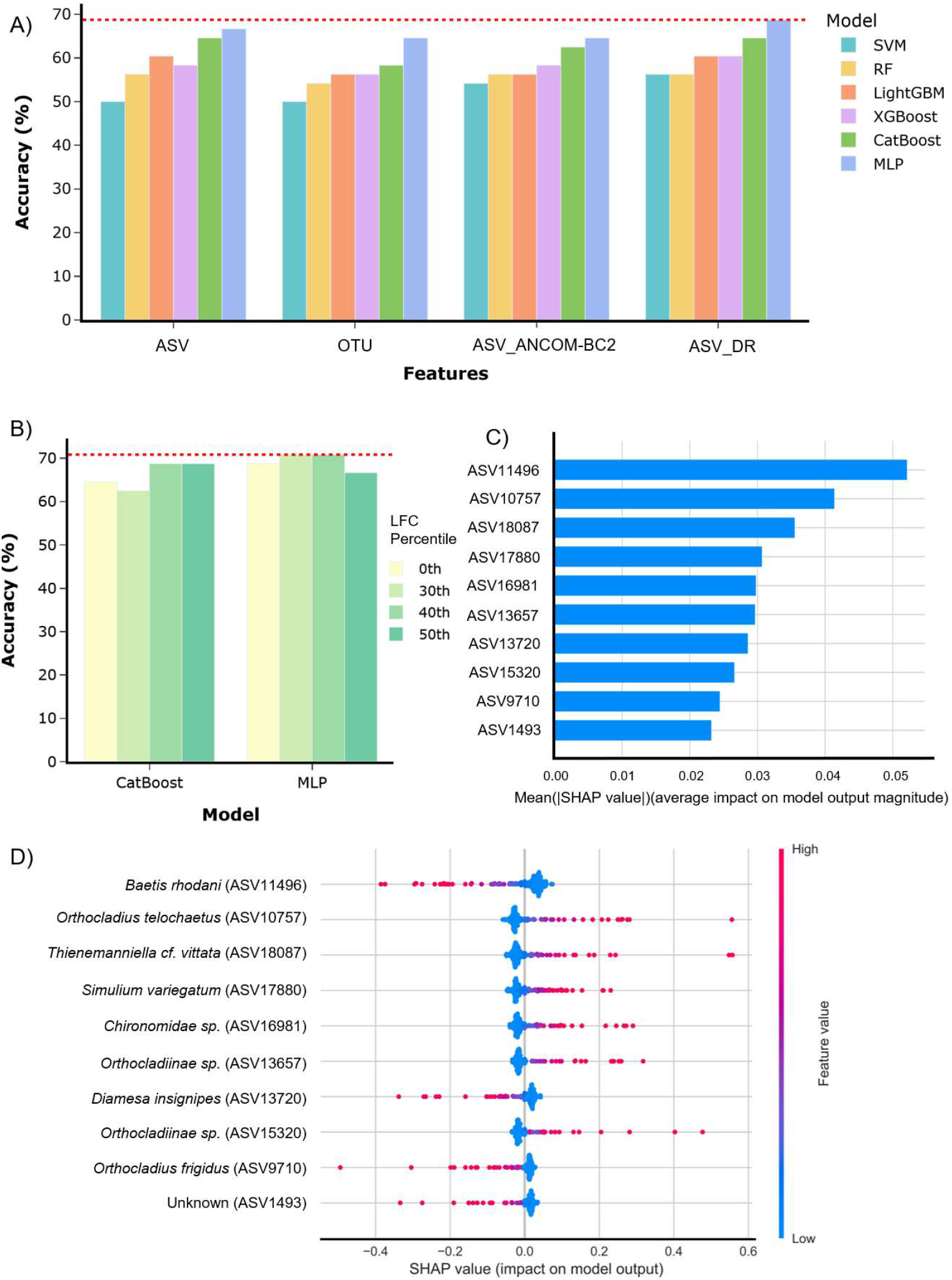
Impact prediction with ML using eDNA data. A) Classification performance of six ML models using ASV, OTU, ASV_ANCOM-BC2 or ASV_DR datasets. Best classification accuracy indicated by red dashed line; B) Effect of log-fold change (LFC) percentile thresholding of ASV_DR dataset on CatBoost and MLP classification performance. Best classification accuracy indicated by red dashed line; C) SHAP bar plot showing feature importance for top 10 ASVs impacting on MLP predictions; D) SHAP beeswarm plot showing how top 10 ASVs affect MLP prediction for the reference class. A feature is positively associated with reference prediction if an increase in value (becoming red) results in an increase in reference predictions (higher positive SHAP values). Taxonomic annotations for ASVs are provided where available.

The potential of further refinement of the ASV_DR dataset by filtering for ASVs that had higher differentials (i.e., larger LFC magnitude and potentially higher signal) to yield performance benefits in ML was examined. The ASV_DR dataset was filtered using minimum thresholds at 30^th^, 40^th^ and 50^th^ percentile of the absolute LFC values, with both MLP and CatBoost models trained on these filtered datasets (Figure 3B). For the MLP model, performance uplifts were observed at 30^th^ and 40^th^ percentile thresholds (both 70.83% accuracy) but degraded at 50^th^ percentile threshold (66.67% accuracy) when compared to the unfiltered dataset (68.75% accuracy) (Figure 3B). In contrast, the CatBoost model performed slightly lower at 30^th^ percentile threshold (62.5% accuracy) but improved at 40^th^ and 50^th^ percentile thresholds (both 68.75% accuracy) when compared to the unfiltered dataset (64.58% accuracy) (Figure 3B). The 40^th^ percentile of LFC was thus selected as the optimum threshold for ASV_DR (hereafter ASV_DR-filtered) as both MLP and CatBoost models exhibited the best performance. The ML pipelines generated by PyCaret for both MLP and CatBoost models showed the same modular workflow adopted for classification (Supporting Information Figure S1). The pipeline consisted of four steps: label encoding, numerical imputing, categorical imputing, and ML classification. In this study, label encoding assigns the value of 0 and 1 to the target classes (impacted=0; reference=1). Both numerical and categorical imputers aim to replace missing values in columns using either the mean or most frequent value but had no impact in this study as there were no missing values in the input table.

SHAP analysis for the best MLP model (70.83% accuracy, based on ASV_DR-filtered) identified key ASVs impacting on prediction (Figure 3C), with taxonomic information available for nine out of the top ten features (Figure 3D). The top three features identified in descending order were *Baetis rhodani* (ASV11496), *Orthocladius telocheatus* (ASV10757) and *Thiememanniella cf. vittata* (ASV18087); the rest had similar contributions to prediction (Figure 3C). The effect of these features on the model predictions for the reference class (class 1) is shown in a beeswarm plot, with the rows corresponding to each feature’s ranking based on mean absolute SHAP value (Figure 3D). A feature’s value is displayed as a dot using a blue (low value) to red (high value) color ramp, with the point position along the x-axis determined by its SHAP value (negative to positive). A feature is said to have a positive impact on reference prediction if an increase in its value (moving towards red) corresponds to an increase in reference class prediction (higher positive SHAP value). Based on this, four out of the top ten features (*Baetis rhodani, Diamese insignipes, Orthocladius frigidus* and an unknown ASV) had a negative impact on reference class prediction with the remaining six (*Orthocladius telochaetus, Thienemanniella cf. vittata, Simulium variegatum, Chironomidae sp.* and two *Orthocladiinae sp.*) showing a positive impact on reference prediction (Figure 3D).

### Impact prediction using combined eDNA and biogeographic data

Classification performance for the six ML models using a combined eDNA and biogeographic features dataset (ASV_DR-filtered + Biogeo) was compared against those obtained using only eDNA (ASV_DR-filtered) (Figure 4A). The results show a marked improvement in performance across all ML models using the combined multimodal dataset (66.67 – 83.33 % accuracy) when compared to those using eDNA data only (56.25 – 70.83% accuracy). The most significant enhancement was observed in the MLP model, coincidentally the best model, with a 12.5% performance uplift when transitioning from eDNA only (70.83% accuracy) to the combined dataset (83.33% accuracy) (Figure 4A). The MLP model also had a significant performance lead over other models (second best CatBoost model had 75% accuracy) with the best scores in precision, recall and F1-score (Supporting Information Table S2). Examination of the MLP pipelines produced by PyCaret showed an increase in network complexity when transitioning from eDNA data (single hidden layer, Figure S1) to combined data (three hidden layers, Figure S2), which may have contributed to the capturing of salient features in the multimodal dataset. The tree-based models (RF, LightGBM, XGBoost and CatBoost) also benefited from the addition of biogeographic features in the combined dataset, with classification accuracies exceeding 72% (72.92 – 75% accuracy). The SVM model had the lowest classification accuracy of 66.67% on the combined dataset, although this still represented a significant 10.42% uplift from using eDNA data only (Figure 4A). Examination of the ML pipelines for both MLP and CatBoost models for the combined dataset showed a similar workflow as that for eDNA previously, except for an additional one-hot encoding step prior to ML classification (Figure 4B, Supporting Information Figure S2). One-hot encoding converts categorical data into a numerical format suitable for ML, by generating binary (0 or 1) columns for each unique category, indicating its presence or absence. The effect of one-hot encoding and label encoding on the combined feature table in this study is illustrated in Figure 4C.

**Figure 4.**
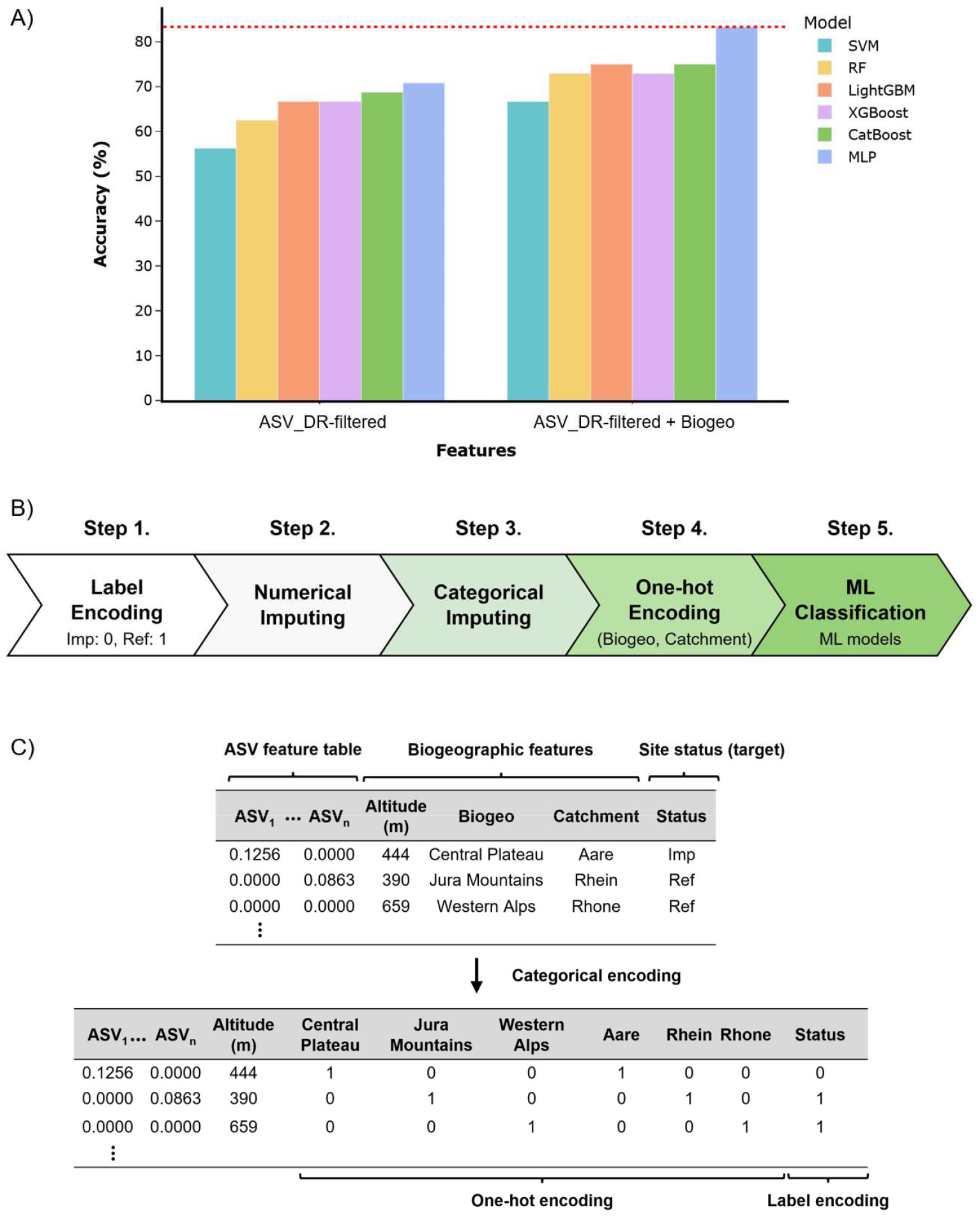
Impact prediction with ML using combined eDNA and biogeographic data. A) Classification performance of six ML models using eDNA only (ASV_DR-filtered) or combined eDNA with biogeographic features (ASV_DR-filtered + Biogeo). The biogeographic features (biogeographic region, altitude, catchment) were merged directly to the ASV feature table. Best classification accuracy indicated by red dashed line; B) PyCaret ML pipeline for impact prediction using combined multimodal dataset; C) Effect of one-hot encoding (present=1, absent=0 for each unique value in a categorical variable) and label encoding (Ref=1, Imp=0 for site status) on the combined input feature table for ML.

SHAP analysis for all six ML models on the combined dataset confirmed that biogeographic features were making significant contributions to model predictions (Supporting Information Figure S3). Except for the SVM model which had altitude as the top feature, all other models had the biogeographic region of Central Plateau as the top feature. Compared to the tree-based models, the MLP model had more biogeographic features (six) within the top ten ranking; the tree-based models had two biogeographic features (Central Plateau and altitude) (Figure S3). These results suggest that the MLP model was able to better utilize the different modalities afforded by the combined dataset. The results also indicate that MLP and tree-based models, at least in this study, were more amenable to mixed data types without being affected by feature scaling, as opposed to the SVM model which was likely affected by the altitude feature’s larger range and magnitude (198 to 1650, as opposed to values between 0 and 1) (Figure S3). Separate testing on the impact of the altitude feature scaled to the range 0 to 1 (MinMaxScaler) or Z-score normalized (StandardScaler) on MLP performance showed that both approaches had a detrimental effect (Supporting Information Figure S4). SHAP plots for the best MLP model showed that three ASVs (*B. rhodani, O. telocheatus* and *S. varigatum*) previously identified as key features for MLP using eDNA data were also present, with the addition of *Anocha sp.* (ASV857) (Figure 5A, 5B). Four features (Biogeo_Central Plateau, altitude, *B. rhodani* and catchment_Aare) had a negative impact on the reference class prediction, with the remaining six features (Biogeo_Jura Mountains, Catchment_Rhein, Biogeo_Northern Alps, *Anocha sp., O. telocheatus* and *S. varigatum*) having a positive impact on reference predictions (Figure 5B). The biomonitoring site distribution map (see Figure 1) showed that most sites within the biogeographic region of Central Plateau were impacted, thus explaining its negative association with the reference class. Similarly, most sites within the biogeographic regions of Jura Mountains and Northern Alps had reference status, supporting the observation that these regions had a positive impact on reference predictions.

**Figure 5.**
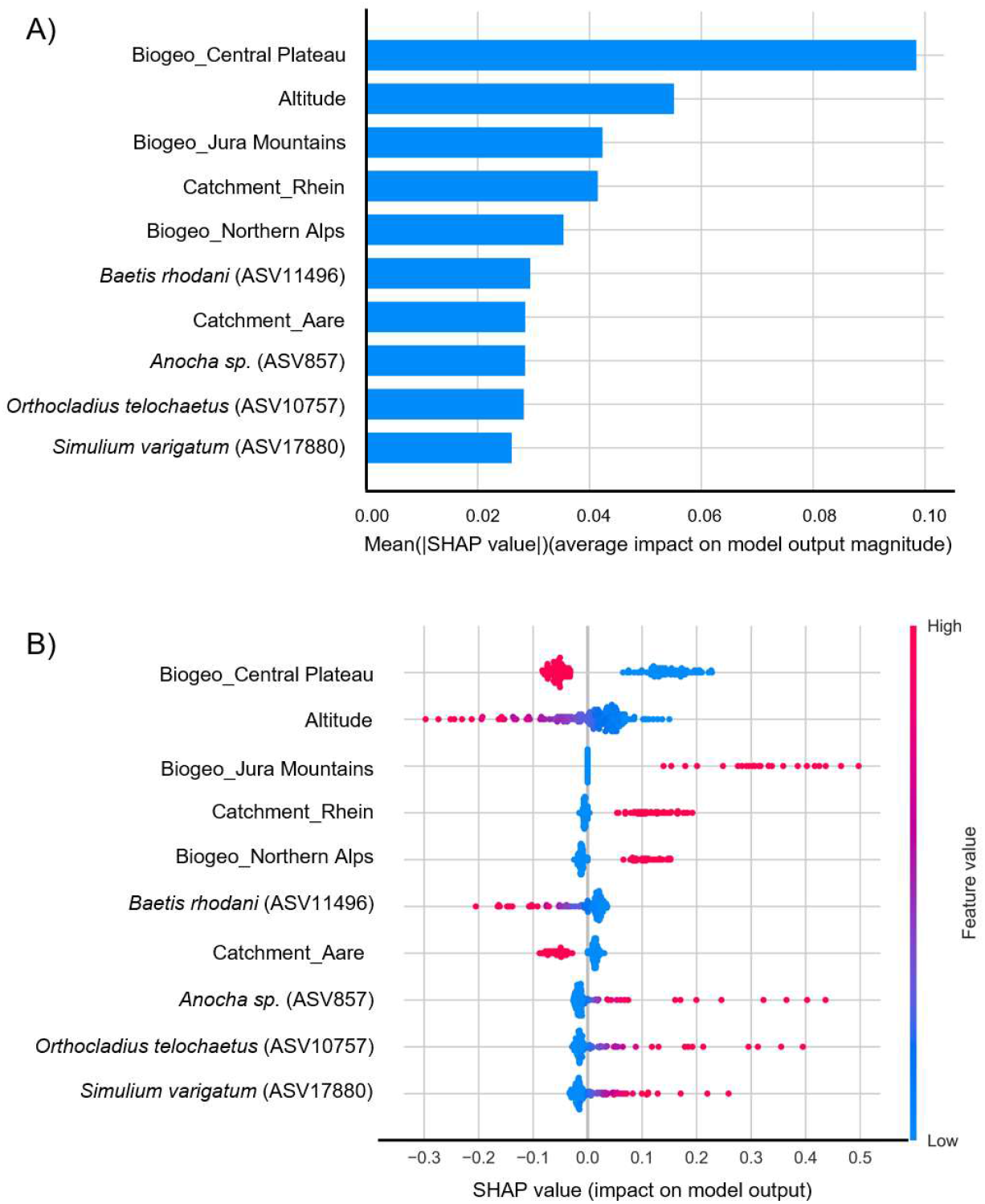
SHAP visualizations for the top 10 features impacting MLP predictions using combined eDNA and biogeographic data. A) Bar plot showing feature importance summarized as mean absolute SHAP value or magnitude; B) Beeswarm plot showing how each feature affects MLP prediction for the reference class. Taxonomic annotations for ASVs are provided where available.

## DISCUSSION

In this study, using impact prediction for Switzerland’s freshwater biomonitoring sites as an example, I introduce a multimodal ML workflow using combined eDNA (ASVs) and biogeographic data to directly infer environmental states. Results showed that across six popular ML models, the multimodal ML approach significantly outperformed ML using only eDNA data. The success of the proposed workflow can be attributed to synergistic components that enhance ML performance by selecting and enriching for high-quality features, and harnessing additional information contained in the combined dataset.

The first step in the proposed workflow aims to select high-quality features from eDNA sequence data and achieve concurrent reduction in data dimensionality in preparation for feature merging with biogeographic data. Comparison of ML performance obtained from four datasets corresponding to the total ASVs (50 – 66.67% accuracy), OTUs (50 – 64.58% accuracy) and differentially abundant ASVs identified by ANCOM-BC2 (54.17 – 64.58% accuracy) and DR (56.25 – 68.75% accuracy) showed that both OTU and DA analysis were effective dimension reduction methods, with DA analyses achieving comparable or better performance than the total ASVs. These results are comparable to the 69.1% classification accuracy achieved by a RF model in a previous study using OTUs for impact prediction across Switzerland’s biomonitoring sites ^8^, although it is noteworthy that the reported result was obtained from a 10-fold cross validated training set (i.e., not from unseen data or test set) with a smaller sample size (64 sites) collected from a single spring season. The construction of de novo OTUs by clustering sequencing reads into a consensus sequence (the OTU) based on an arbitrary sequence similarity threshold (usually 97%) is a customary step in eDNA and microbiome studies ^25, 39^. Compared to ASVs which are exact sequences, de novo OTUs offer lower taxonomic resolution and are dataset-dependent with varied outcomes depending on the chosen similarity threshold ^25^. For enhanced taxonomic resolution and reproducibility, ASVs are preferred but are not generally used in large-scale studies due to high computational costs associated with data processing.

In this study, DA analysis achieved very significant reductions of 94 – 99% in ASV data size whilst preserving key features impacting on model predictions, making it an ideal approach for feature selection and dimension reduction in ML, and possibly as a strategy to address large ASV datasets in general. Compared to DR, ANCOM-BC2 imposed a more stringent selection on ASVs, although all significant ASVs identified were also present in those selected by DR. ANCOM-BC2 is an extension of ANCOM-BC (Analysis of Compositions of Microbiomes with Bias Correction) ^40^, with enhancements for multigroup analysis and repeated measures ^23^. Both ANCOM-BC and ANCOM-BC2 identify differentially abundant taxa modelled using a linear regression framework via statistical testing, with ANCOM-BC2 implementing a stricter selection regime with improved bias correction, variance regularization and sensitivity analysis leading to much lower false discovery rates (FDR) ^23^. In contrast, DR adopts a ranking approach that ranks taxa/features based on their relative differentials (log ratio between relative abundances) estimated using a multinomial regression ^24^. Studies have shown that both ANCOM-BC and DR can have large overlaps (50 – 80%) of differentially abundant taxa/features ^24, 28^, with both methods generally considered to be complementary. However, it is known that different DA methods can produce drastically different outcomes depending on the dataset ^41^, thus it may be beneficial to adopt a consensus approach based on multiple DA approaches in future to identify significant ASVs and ensure robust biological interpretations. Initial testing in this study showed that the ASV_DR dataset yielded the best ML performance with MLP (68.75% accuracy), with further improvements (70.83% accuracy) obtained via LFC percentile thresholding. The optimum LFC threshold should be determined empirically for different datasets.

The optimal filtered ASV feature table from DR was merged directly with biogeographic data (biogeographic region, altitude and catchment area) without further data manipulation (e.g., feature transformation or scaling) to create a combined multimodal dataset. This step was streamlined in large due to the use of an AutoML framework, PyCaret, that automates data preprocessing and encoding of categorical variables prior to model training. Feature scaling which standardizes the range of features was disabled as this operation is dataset-dependent and may have detrimental effects on ML performance (e.g., as in this study for MLP, Figure S4). Although tree-based models such as RF, LightGBM, XGBoost and CatBoost are tolerant to varied feature magnitudes, distance-based models such as SVM (which aims to achieve maximum distance/separation between class boundaries) and neural networks such as MLP (constructed from layers of densely connected nodes, i.e., neurons) are more sensitive to feature scaling ^42^. Nevertheless, significant performance uplifts were observed across all six ML models when transitioning from eDNA data (56.25 – 70.83% accuracy) to the combined dataset (66.67 – 83.33% accuracy), providing strong support for a multimodal ML approach. The best classification performance was recorded by the MLP model (83.33% accuracy), followed by the tree-based models (72.92 – 75% accuracy) and lastly the SVM model (66.67% accuracy). The results suggest that the MLP model may be more efficient in utilizing information present in the combined dataset, in part due to its ability to vary model complexity (i.e., increasing network layers) to accommodate additional modalities. Future ML studies may consider deep neural networks, such as convolutional neural networks (CNNs) which have found wide application and success in the genomic sciences ^43, 44^.

The tree-based models also performed well across ML experiments in this study, with CatBoost consistently finishing as second best. Of these, LightGBM, XGBoost and CatBoost are variations of the gradient boosted decisions trees (GBDTs) algorithm which combines multiple weak learners (decision trees) to create a strong learner (i.e., ensemble learning) in a sequential and iterative manner with each subsequent tree correcting the errors made by previous trees (a process called boosting) ^33^. In contrast, the RF algorithm, which is also an ensemble ML method, constructs distinct decision trees in parallel using a random subset of features and training data (a process called bagging) ^45^. Results showed that the trio of GBDTs overall outperformed RF and SVM across ML experiments, supporting these models as important inclusions in future ML studies. Studies have shown that GBDTs, specifically XGBoost and CatBoost, were superior to other ML models including deep learning frameworks for classification and regression problems using tabular data ^46, 47^. Both XGBoost and CatBoost are highly scalable and tolerant of varied feature scales, making them potentially the better option when dealing with very large and complex multimodal tabular datasets compared to neural networks or models sensitive to feature scaling.

The final step in the proposed multimodal ML workflow is model explanation which uses explainable artificial intelligence (XAI) techniques to understand ML model behavior and output. A feature-based XAI method, SHAP, was implemented in this study as it is model agnostic and widely used in ML studies ^34, 35^. For the best model (MLP) using eDNA data, SHAP analysis identified *Baetis rhodani,* one of the most widespread mayfly species, as the top ranked ASV feature with a negative impact on reference prediction (i.e., increased relative abundance is associated with the impact class). A recent study showed that *Baetis rhodani* had a higher abundance in urban areas compared to natural environments, with a positive correlation with concrete cover ^48^. It has been shown that *Baetis rhodani* is tolerant against potential urban effects, such as heavy metal pollution ^49^ and has a broad substrate range with a preference for high water velocity ^50^, which is typical of urban streams. A review of existing literature largely supports the top-ranked ASV features (where species identity is known) as having biological or ecological significance, with some used as bioindicator species for water quality or environmental pollution (Supporting Information Table S3). The success of SHAP analysis in uncovering features with biological significance can be largely attributed to the use of DA analysis for feature selection at the start in the workflow, supporting the combination of DA analysis and SHAP as a powerful tool for discovering ecologically significant taxa in future eDNA ML studies.

With the combined multimodal dataset, SHAP analysis showed that biogeographic features contributed significantly to ML model predictions, concomitant with the increase in classification performance across ML models. For the best model (MLP), six out of the top 10 features were derived from biogeographic data, with the remaining four from eDNA (ASVs). Of the four significant ASVs, three were also identified previously in MLP using eDNA, with the addition of *Anthoca sp.* which had a positive association with the reference prediction. *Anthoca sp.,* a type of stonefly, has been proposed as a bioindicator for good to very good water quality, as it is found in clear streams and sensitive/intolerant to pollutants ^51^. The biogeographic region of Central Plateau was the top ranked feature followed by altitude, with both features negatively associated with the reference class. The Central Plateau covers 30% of Switzerland but is the most densely populated region (2/3 of population), with close to 50% of land used for agriculture, while forests cover around 24% ^52^. Thus, it is no surprise that majority of the impacted sites were located within the Central Plateau. However, the impact of altitude on model prediction was less obvious and may be confounded by interaction with other factors such as the dominant Central Plateau feature, coincidentally also the most frequent region found in approximately half of the training samples. This highlights the potential weakness of the SHAP method, in that it assumes all features/variables are independent, which may not always be true ^53, 54^. This concern may be mitigated by removing closely correlated features prior to ML, or by implementing modified kernel SHAP methods that account for feature dependence ^54^. Alternately, post-SHAP methods such as Normalized Movement Rate (NMR) ^55^ and Modified Index Position (MIP) ^56^ which adjust for collinearity between features may also be considered. Although SHAP values are model agnostic, they are nonetheless model-dependent, meaning different features and ranking could result from different models ^36^. Where there are multiple similarly performing models, careful consideration of top-ranked features common across models is required as SHAP values are not directly comparable between models. However, it is possible to discern how each model weighs the different features by considering the shape of the different SHAP plots ^57^, including force-plots ^35^ which depict how individual features affect a specific prediction for a given data point.

The multimodal ML workflow introduced in this study is applicable to biomonitoring programs across large spatiotemporal scales, with the adoption of ASVs as the standard eDNA analytical unit making it transferable across programs at the national and international levels where standardized biogeographic inputs and representative training data are available or possible. For smaller biomonitoring programs, specifically longitudinal studies with few targeted sites, increasing sampling frequency or number and treating each sample as individual data points may alleviate the training data requirement for ML. The sustained explosion in eDNA data encouraged by advancements in high-throughput sequencing and standardized eDNA collection protocols may mitigate ML training data requirements moving forward. The proposed workflow can also be adapted for multiclass or regression problems, capitalizing on DA analysis methods (e.g., ANCOM-BC2 and DR) that can conduct multigroup testing and AutoML frameworks that are readily configured for classification or regression tasks. Perhaps the most exciting aspect of the proposed workflow is the limitless possibilities afforded by the combination of eDNA data with modalities extending beyond biogeographic data in ML, such as those derived from omics (e.g., proteomics and metabolomics) or remote sensing (e.g., spectral, 3D structural, thermal and radar) for a holistic understanding of ecosystem health and functioning.

## AUTHOR INFORMATION

### Present Addresses

†If an author’s address is different than the one given in the affiliation line, this information may be included here.

### Author Contributions

Joshua C.O. Koh designed the study, analyzed the data, prepared the figures and wrote the manuscript.

### Notes

All relevant source codes and datasets required to reproduce results in this study, including the best MLP models reported are available at https://github.com/OSS-DOTS/eDNA-Multimodal-ML.

### Supporting Information

Supporting information: taxonomic assignment of top-ranked ASVs from MLP (Table S1), PyCaret ML pipelines (Figure S1 and S2), SHAP bar plots for ML models using combined multimodal dataset (Figure S3), classification report for multimodal ML models (Table S2), ecological significance of top ranked ASVs from MLP (Table S3) and effect of feature scaling on MLP model are provided in the supporting information.pdf file.

## ACKNOWLEDGMENT

I wish to thank Dr Che Doering for his helpful comments on the manuscript.

## REFERENCES

(1) Diaz, S.; Settele, J.; Brondizio, E. S.; Ngo, H. T.; Agard, J.; Arneth, A.; Balvanera, P.; Brauman, K. A.; Butchart, S. H. M.; Chan, K. M. A.; et al. Pervasive human-driven decline of life on Earth points to the need for transformative change. Science 2019, 366 (6471). DOI: 10.1126/science.aax3100 From NLM Medline.

(2) Global assessment report on biodiversity and ecosystem services of the Intergovernmental Science-Policy Platform on Biodiversity and Ecosystem Services; 2019. DOI: 10.5281/zenodo.6417333.

(3) Isbell, F.; Balvanera, P.; Mori, A. S.; He, J.-S.; Bullock, J. M.; Regmi, G. R.; Seabloom, E. W.; Ferrier, S.; Sala, O. E.; Guerrero-Ramírez, N. R.; et al. Expert perspectives on global biodiversity loss and its drivers and impacts on people. Frontiers in Ecology and the Environment 2023, 21 (2), 94–103. DOI: 10.1002/fee.2536.

(4) Cordier, T.; Alonso-Saez, L.; Apotheloz-Perret-Gentil, L.; Aylagas, E.; Bohan, D. A.; Bouchez, A.; Chariton, A.; Creer, S.; Fruhe, L.; Keck, F.; et al. Ecosystems monitoring powered by environmental genomics: A review of current strategies with an implementation roadmap. Mol Ecol 2021, 30 (13), 2937–2958. DOI: 10.1111/mec.15472 From NLM Medline.

(5) Pawlowski, J.; Bonin, A.; Boyer, F.; Cordier, T.; Taberlet, P. Environmental DNA for biomonitoring. Mol Ecol 2021, 30 (13), 2931–2936. DOI: 10.1111/mec.16023 From NLM Medline.

(6) Leese, F.; Bouchez, A.; Abarenkov, K.; Altermatt, F.; Borja, Á.; Bruce, K.; Ekrem, T.; Čiampor, F.; Čiamporová-Zaťovičová, Z.; Costa, F. O.; et al. Chapter Two - Why We Need Sustainable Networks Bridging Countries, Disciplines, Cultures and Generations for Aquatic Biomonitoring 2.0: A Perspective Derived From the DNAqua-Net COST Action. In Advances in Ecological Research, Bohan, D. A., Dumbrell, A. J., Woodward, G., Jackson, M. Eds.; Vol. 58; Academic Press, 2018; pp 63–99.

(7) Keck, F.; Blackman, R. C.; Bossart, R.; Brantschen, J.; Couton, M.; Hurlemann, S.; Kirschner, D.; Locher, N.; Zhang, H.; Altermatt, F. Meta-analysis shows both congruence and complementarity of DNA and eDNA metabarcoding to traditional methods for biological community assessment. Mol Ecol 2022, 31 (6), 1820–1835. DOI: 10.1111/mec.16364 From NLM Medline.

(8) Keck, F.; Brantschen, J.; Altermatt, F. A combination of machine-learning and eDNA reveals the genetic signature of environmental change at the landscape levels. Mol Ecol 2023, 32 (17), 4791–4800. DOI: 10.1111/mec.17073 From NLM Medline.

(9) Apothéloz-Perret-Gentil, L.; Cordonier, A.; Straub, F.; Iseli, J.; Esling, P.; Pawlowski, J. Taxonomy-free molecular diatom index for high-throughput eDNA biomonitoring. Molecular Ecology Resources 2017, 17 (6), 1231–1242. DOI: 10.1111/1755-0998.12668.

(10) Feio, M. J.; Serra, S. R. Q.; Mortágua, A.; Bouchez, A.; Rimet, F.; Vasselon, V.; Almeida, S. F. P. A taxonomy-free approach based on machine learning to assess the quality of rivers with diatoms. Science of The Total Environment 2020, 722, 137900. DOI: 10.1016/j.scitotenv.2020.137900.

(11) Cordier, T.; Esling, P.; Lejzerowicz, F.; Visco, J.; Ouadahi, A.; Martins, C.; Cedhagen, T.; Pawlowski, J. Predicting the Ecological Quality Status of Marine Environments from eDNA Metabarcoding Data Using Supervised Machine Learning. Environ Sci Technol 2017, 51 (16), 9118–9126. DOI: 10.1021/acs.est.7b01518 From NLM Medline.

(12) Fruhe, L.; Cordier, T.; Dully, V.; Breiner, H. W.; Lentendu, G.; Pawlowski, J.; Martins, C.; Wilding, T. A.; Stoeck, T. Supervised machine learning is superior to indicator value inference in monitoring the environmental impacts of salmon aquaculture using eDNA metabarcodes. Mol Ecol 2021, 30 (13), 2988–3006. DOI: 10.1111/mec.15434 From NLM Medline.

(13) Brantschen, J.; Blackman, R. C.; Walser, J. C.; Altermatt, F. Environmental DNA gives comparable results to morphology-based indices of macroinvertebrates in a large-scale ecological assessment. PLoS One 2021, 16 (9), e0257510. DOI: 10.1371/journal.pone.0257510 From NLM Medline.

(14) Brown, C. J.; Saunders, M. I.; Possingham, H. P.; Richardson, A. J. Managing for interactions between local and global stressors of ecosystems. PLoS One 2013, 8 (6), e65765. DOI: 10.1371/journal.pone.0065765 From NLM Medline.

(15) Geary, W. L.; Bode, M.; Doherty, T. S.; Fulton, E. A.; Nimmo, D. G.; Tulloch, A. I. T.; Tulloch, V. J. D.; Ritchie, E. G. A guide to ecosystem models and their environmental applications. Nat Ecol Evol 2020, 4 (11), 1459–1471. DOI: 10.1038/s41559-020-01298-8 From NLM Medline.

(16) Cordier, T.; Forster, D.; Dufresne, Y.; Martins, C. I. M.; Stoeck, T.; Pawlowski, J. Supervised machine learning outperforms taxonomy-based environmental DNA metabarcoding applied to biomonitoring. Mol Ecol Resour 2018, 18 (6), 1381–1391. DOI: 10.1111/1755-0998.12926 From NLM Medline.

(17) NAWA nationale Beobachtung Oberflächengewässerqualität. – Bundesamt für Umwelt, Bern.; BAFU, 2013.

(18) Loran, C.; Munteanu, C.; Verburg, P. H.; Schmatz, D. R.; Bürgi, M.; Zimmermann, N. E. Long-term change in drivers of forest cover expansion: an analysis for Switzerland (1850-2000). Regional Environmental Change 2017, 17 (8), 2223–2235. DOI: 10.1007/s10113-017-1148-y.

(19) Vamos, E. E.; Elbrecht, V.; Leese, F. Short COI markers for freshwater macroinvertebrate metabarcoding. Metabarcoding and Metagenomics 2017, 1. DOI: 10.3897/mbmg.1.14625.

(20) Leese, F.; Sander, M.; Buchner, D.; Elbrecht, V.; Haase, P.; Zizka, V. M. A. Improved freshwater macroinvertebrate detection from environmental DNA through minimized nontarget amplification. Environmental DNA 2021, 3 (1), 261–276. DOI: 10.1002/edn3.177.

(21) Callahan, B. J.; McMurdie, P. J.; Rosen, M. J.; Han, A. W.; Johnson, A. J. A.; Holmes, S. P. DADA2: High-resolution sample inference from Illumina amplicon data. Nature Methods 2016, 13 (7), 581–583. DOI: 10.1038/nmeth.3869.

(22) Rognes, T.; Flouri, T.; Nichols, B.; Quince, C.; Mahe, F. VSEARCH: a versatile open source tool for metagenomics. PeerJ 2016, 4, e2584. DOI: 10.7717/peerj.2584 From NLM PubMed-not-MEDLINE.

(23) Lin, H.; Peddada, S. D. Multigroup analysis of compositions of microbiomes with covariate adjustments and repeated measures. Nat Methods 2024, 21 (1), 83–91. DOI: 10.1038/s41592-023-02092-7 From NLM Medline.

(24) Morton, J. T.; Marotz, C.; Washburne, A.; Silverman, J.; Zaramela, L. S.; Edlund, A.; Zengler, K.; Knight, R. Establishing microbial composition measurement standards with reference frames. Nat Commun 2019, 10 (1), 2719. DOI: 10.1038/s41467-019-10656-5 From NLM Medline.

(25) Callahan, B. J.; McMurdie, P. J.; Holmes, S. P. Exact sequence variants should replace operational taxonomic units in marker-gene data analysis. The ISME Journal 2017, 11 (12), 2639–2643. DOI: 10.1038/ismej.2017.119.

(26) Wilhelm, R. C.; van Es, H. M.; Buckley, D. H. Predicting measures of soil health using the microbiome and supervised machine learning. Soil Biology and Biochemistry 2022, 164, 108472. DOI: 10.1016/j.soilbio.2021.108472.

(27) Ghannam, R. B.; Techtmann, S. M. Machine learning applications in microbial ecology, human microbiome studies, and environmental monitoring. Comput Struct Biotechnol J 2021, 19, 1092–1107. DOI: 10.1016/j.csbj.2021.01.028 From NLM PubMed-not-MEDLINE.

(28) Lin, H.; Peddada, S. D. Analysis of microbial compositions: a review of normalization and differential abundance analysis. npj Biofilms and Microbiomes 2020, 6 (1), 60. DOI: 10.1038/s41522-020-00160-w.

(29) Pedregosa, F.; Varoquaux, G.; Gramfort, A.; Michel, V.; Thirion, B.; Grisel, O.; Blondel, M.; Prettenhofer, P.; Weiss, R.; Dubourg, V. Scikit-learn: Machine learning in Python. the Journal of machine Learning research 2011, 12, 2825–2830.

(30) Waring, J.; Lindvall, C.; Umeton, R. Automated machine learning: Review of the state-of-the-art and opportunities for healthcare. Artificial Intelligence in Medicine 2020, 104, 101822. DOI: 10.1016/j.artmed.2020.101822.

(31) PyCaret: An open source, low-code machine learning library in Python; 2020. https://www.pycaret.org (accessed 26-04-2025).

(32) Dhall, D.; Kaur, R.; Juneja, M. Machine Learning: A Review of the Algorithms and Its Applications. Cham, 2020; Springer International Publishing: pp 47–63.

(33) Bentéjac, C.; Csörgő, A.; Martínez-Muñoz, G. A comparative analysis of gradient boosting algorithms. Artificial Intelligence Review 2021, 54 (3), 1937–1967. DOI: 10.1007/s10462-020-09896-5.

(34) Hassija, V.; Chamola, V.; Mahapatra, A.; Singal, A.; Goel, D.; Huang, K.; Scardapane, S.; Spinelli, I.; Mahmud, M.; Hussain, A. Interpreting Black-Box Models: A Review on Explainable Artificial Intelligence. Cognitive Computation 2024, 16 (1), 45–74. DOI: 10.1007/s12559-023-10179-8.

(35) Lundberg, S. M.; Lee, S.-I. A unified approach to interpreting model predictions. In Proceedings of the 31st International Conference on Neural Information Processing Systems, Long Beach, California, USA; 2017.

(36) Ponce-Bobadilla, A. V.; Schmitt, V.; Maier, C. S.; Mensing, S.; Stodtmann, S. Practical guide to SHAP analysis: Explaining supervised machine learning model predictions in drug development. Clin Transl Sci 2024, 17 (11), e70056. DOI: 10.1111/cts.70056 From NLM.

(37) Raschka, S. Model evaluation, model selection, and algorithm selection in machine learning. arXiv preprint 1811.12808 2018.

(38) Akiba, T.; Sano, S.; Yanase, T.; Ohta, T.; Koyama, M. Optuna: A next-generation hyperparameter optimization framework. In Proceedings of the 25th ACM SIGKDD international conference on knowledge discovery & data mining, 2019; pp 2623–2631.

(39) Knight, R.; Vrbanac, A.; Taylor, B. C.; Aksenov, A.; Callewaert, C.; Debelius, J.; Gonzalez, A.; Kosciolek, T.; McCall, L.-I.; McDonald, D. Best practices for analysing microbiomes. Nature Reviews Microbiology 2018, 16 (7), 410–422.

(40) Lin, H.; Peddada, S. D. Analysis of compositions of microbiomes with bias correction. Nature Communications 2020, 11 (1), 3514. DOI: 10.1038/s41467-020-17041-7.

(41) Nearing, J. T.; Douglas, G. M.; Hayes, M. G.; MacDonald, J.; Desai, D. K.; Allward, N.; Jones, C. M. A.; Wright, R. J.; Dhanani, A. S.; Comeau, A. M.; et al. Microbiome differential abundance methods produce different results across 38 datasets. Nature Communications 2022, 13 (1), 342. DOI: 10.1038/s41467-022-28034-z.

(42) de Amorim, L. B. V.; Cavalcanti, G. D. C.; Cruz, R. M. O. The choice of scaling technique matters for classification performance. Applied Soft Computing 2023, 133, 109924. DOI: 10.1016/j.asoc.2022.109924.

(43) Eraslan, G.; Avsec, Ž.; Gagneur, J.; Theis, F. J. Deep learning: new computational modelling techniques for genomics. Nature Reviews Genetics 2019, 20 (7), 389–403. DOI: 10.1038/s41576-019-0122-6.

(44) Montesinos-López, O. A.; Montesinos-López, A.; Pérez-Rodríguez, P.; Barrón-López, J. A.; Martini, J. W. R.; Fajardo-Flores, S. B.; Gaytan-Lugo, L. S.; Santana-Mancilla, P. C.; Crossa, J. A review of deep learning applications for genomic selection. BMC Genomics 2021, 22 (1), 19. DOI: 10.1186/s12864-020-07319-x.

(45) Biau, G.; Scornet, E. A random forest guided tour. Test 2016, 25 (2), 197–227.

(46) Prokhorenkova, L.; Gusev, G.; Vorobev, A.; Dorogush, A. V.; Gulin, A. CatBoost: unbiased boosting with categorical features. In 2018.

(47) Shwartz-Ziv, R.; Armon, A. Tabular data: Deep learning is not all you need. Information Fusion 2022, 81, 84–90. DOI: 10.1016/j.inffus.2021.11.011.

(48) Bohus, A.; Gál, B.; Barta, B.; Szivák, I.; Karádi-Kovács, K.; Boda, P.; Padisák, J.; Schmera, D. Effects of urbanization-induced local alterations on the diversity and assemblage structure of macroinvertebrates in low-order streams. Hydrobiologia 2023, 850 (4), 881–899. DOI: 10.1007/s10750-022-05130-1.

(49) G., R.; W., S. Heavy Metal Accumulation by Baetis rhodani and Macrobenthic Community Structure in Running Waters of the N’ Harz Mountains (Lower Saxony/FRG) (Ephemeroptera: Baetidae). Entomologia Generalis 1991, 16 (1), 31–37. DOI: 10.1127/entom.gen/16/1991/31.

(50) Vilenica, M.; Brigić, A.; Sartori, M.; Mihaljević, Z. Microhabitat selection and distribution of functional feeding groups of mayfly larvae (Ephemeroptera) in lotic karst habitats. Knowl. Manag. Aquat. Ecosyst. 2018, (419), 17.

(51) Xu, M.; Wang, Z.; Duan, X.; Pan, B. Effects of pollution on macroinvertebrates and water quality bio-assessment. Hydrobiologia 2014, 729 (1), 247–259. DOI: 10.1007/s10750-013-1504-y.

(52) Swiss Plateau. Swiss federal authorities, 2021. https://www.aboutswitzerland.eda.admin.ch/en/swiss-plateau (accessed 27/04/2025).

(53) Salih, A. M.; Raisi-Estabragh, Z.; Galazzo, I. B.; Radeva, P.; Petersen, S. E.; Lekadir, K.; Menegaz, G. A Perspective on Explainable Artificial Intelligence Methods: SHAP and LIME. Advanced Intelligent Systems 2025, 7 (1), 2400304. DOI: 10.1002/aisy.202400304.

(54) Aas, K.; Jullum, M.; Løland, A. Explaining individual predictions when features are dependent: More accurate approximations to Shapley values. Artificial Intelligence 2021, 298, 103502. DOI: 10.1016/j.artint.2021.103502.

(55) Salih, A.; Galazzo, I. B.; Cruciani, F.; Brusini, L.; Radeva, P. Investigating Explainable Artificial Intelligence for MRI-based Classification of Dementia: a New Stability Criterion for Explainable Methods. In 2022 IEEE International Conference on Image Processing (ICIP), 16-19 Oct. 2022, 2022; pp 4003–4007. DOI: 10.1109/ICIP46576.2022.9897253.

(56) Salih, A. M.; Galazzo, I. B.; Raisi-Estabragh, Z.; Petersen, S. E.; Menegaz, G.; Radeva, P. Characterizing the Contribution of Dependent Features in XAI Methods. IEEE J Biomed Health Inform 2024, 28 (11), 6466–6473. DOI: 10.1109/jbhi.2024.3395289 From NLM.

(57) Stenwig, E.; Salvi, G.; Rossi, P. S.; Skjærvold, N. K. Comparative analysis of explainable machine learning prediction models for hospital mortality. BMC Med Res Methodol 2022, 22 (1), 53. DOI: 10.1186/s12874-022-01540-w From NLM.

