## Supporting Information for "Explainable Multimodal Machine Learning Using Combined Environmental DNA and Biogeographic Features for Ecosystem Biomonitoring"

### Table S1. Taxonomic assignment of top-ranked ASV features for MLP model using eDNA and combined multimodal dataset

| ASV ID | DNA Sequence | *Keck *et al.* 2023 | | | NCBI-BLASTN | | | Species assigned |
|---|---|---|---|---|---|---|---|---|
| | | Species | Genus confidence (%) | Species confidence (%) | Species | Query Cover (%) | Identities (%) | |
| ASV11496 | TCTCGCCGCAAATATCGCCCACGGAGGGTCTTCGGTTGATTT CGCAATTTTTTCTTTACACTTGGCTGGTATTTCTTCGATTTTAG GTGCAGTTAATTTTATTACTACAGTTGTTAATATGCGTAGCCC TGGAATAACCCTA | *Baetis rhodani* | 95.20 | 95.20 | *Baetis rhodani* | 100.00 | 100.00 | *Baetis rhodani* |
| ASV10757 | TCTATCTTCAGGAATTGCTCACGCCGGAGCATCCGTAGATTT AGCTATTTTTTCCTTACATTTAGCCGGTATTTCTTCTATTTTAG GAGCTGTTAATTTTATTACAACTGTAATTAATATACGATCAAAT GGTATTACTCTT | *Simulium latipes* | 11.43 | 3.93 | *Orthocladius telochaetus* | 99.00 | 93.57 | *Orthocladius telochaetus* |
| ASV18087 | TTTATCGTCTAATATTGCTCATGCAGGTGCTTCAGTAGATTTA GCAATTTTTTCTTTACATTTAGCCGGAATCTCTTCAATTTTAGG GGCTGTAAATTTTATTACAACTGTAATTAATATACGGTCTGAA GGAATTTCATTA | *Tokunagaia kibunensis* | 6.02 | 6.02 | *Thienemanniella cf. vittata* | 100.00 | 92.96 | *Thienemanniella cf. vittata* |
| ASV17880 | TTTATCCTCTGGAATTGCTCATGCCGGAGCTTCCGTTGATTTA GCTATTTTTTCTTTACATTTAGCCGGAATTTCCTCTATTTTAGG AGCTGTAAATTTTATTACAACTATTATTAATATACGATCAAATG GAATTACTTTT | *Simulium variegatum* | 97.52 | 33.39 | *Simulium variegatum* | 100.00 | 100.00 | *Simulium variegatum* |
| ASV16981 | TTTATCATCAAGAATTGCCCATAGAGGAGCATCTGTTGATTTA GCCATTTTTTCTTTACATCTTGCAGGAATTTCTTCAATTTTAGG TTCAGTAAATTTCATTACTACTGCAATTAATATACGATCAACAG GAATTACTCTC | *Dicrotendipes lobiger* | 3.81 | 3.81 | *Chironomidae sp.* | 100.00 | 100.00 | *Chironomidae sp.* |
| ASV13657 | TCTTTCTTCAGGAATCGCACATGCTGGGGCGTCTGTAGATCT AGCCATTTTTTCTTTGCATCTAGCAGGAATTTCCTCTATTTTAG GAGCAGTAAACTTTATTACAACTGTGATTAATATGCGATCTGA CGGTATTACCCTA | *Eukiefferiella minor* | 7.24 | 7.24 | *Orthocladiinae sp.* | 97.00 | 88.41 | *Orthocladiinae sp.* |
| ASV13720 | TCTTTCTTCAGGAATCGCTCATGCTGGAGCATCGGTTGATTTA GCAATTTTTTCTTTACATTTAGCTGGGATTTCTTCTATTTTAGG AGCAGTAAATTTTATTACAACAGTAATTAATATGCGTTCTAGT GGAATTACTTTA | *Diamesa insignipes* | 98.88 | 98.88 | *Diamesa insignipes* | 100.00 | 100.00 | *Diamesa insignipes* |
| ASV15320 | TCTTTCTTCTGGAATTGCTCATGCTGGGGCTTCTGTTGATTTA GCTATTTTTTCTTTACATTTAGCAGGTATTTCTTCTATTTTAGG AGCTGTAAATTTTATTACTACAGTTATTAATATACGATCAAATG GTATTACATTA | *Orthocladius glabripennis* | 13.40 | 6.07 | *Orthocladiinae sp.* | 100.00 | 95.77 | *Orthocladiinae sp.* |
| ASV9710 | GTTATCTTCTGGTATTGCACATGCGGGGGCTTCTGTTGATTTA GCTATTTTTTCTTTACATTTAGCAGGAATTTCTTCTATTTTAGG GGCTGTAAATTTTATTACAACTGTAATTAATATGCGATCAAAT GGAATTACTTTA | *Euryhapsis cilium* | 5.42 | 5.42 | *Orthocladius frigidus* | 99.00 | 95.00 | *Orthocladius frigidus* |
| ASV1493 | ACTTTCAGCAGGATTAGCTCATTCTGGACCAAGTGTTGATATG GCAATATTTAGCTTACACATAGCTGGTTTGTCTTCTCTATTTG GTGCAATAAATTTTATAACTACAATAATAAACATGCGTGTTCC AGGAATGTACTGG | *Strigea indet. species* | 1.46 | 1.46 | *Lissodendoryx sp.* | 55.00 | 84.62 | *Unknown* |
| ASV857 | ACTATCTTCTGGAATTGCTCATACTGGGGCATCAGTAGATTTA GCTATTTTTTCACTACATTTAGCGGGAATTTCATCAATTTTAGG AGCAGTAAATTTTATTACCACTGTAATTAATATACGATCAGCA GGAATTACCTTT | *Propsilocerus akamusi* | 6.06 | 6.06 | *Antocha sp.* | 100.00 | 95.77 | *Antocha sp.* |

*Keck, F.; Brantschen, J.; Altermatt, F. A combination of machine-learning and eDNA reveals the genetic signature of environmental change at the landscape levels. Mol Ecol 2023, 32 (17), 4791-4800.

A) **MLP pipeline: classification accuracy 70.83%**

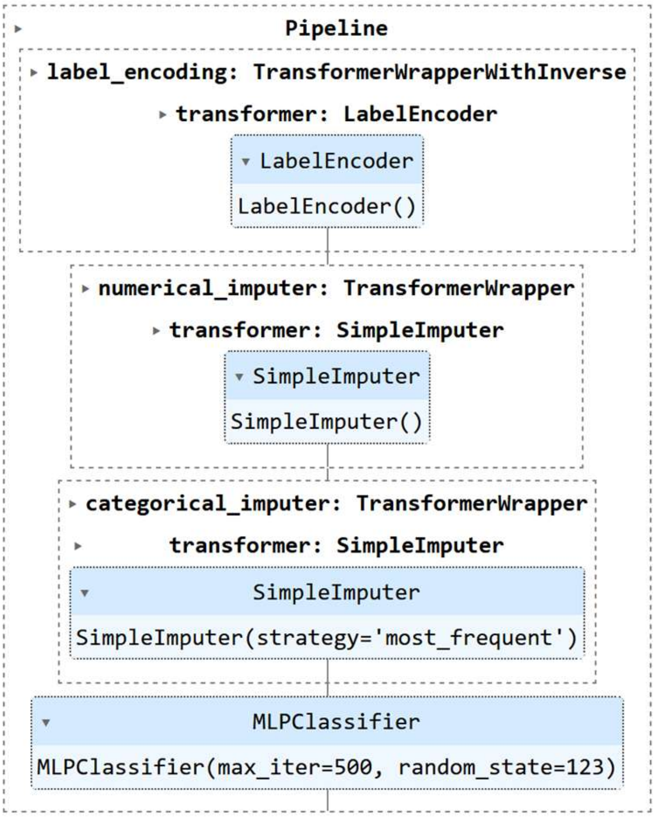


B) **CatBoost pipeline: classification accuracy 68.75%**

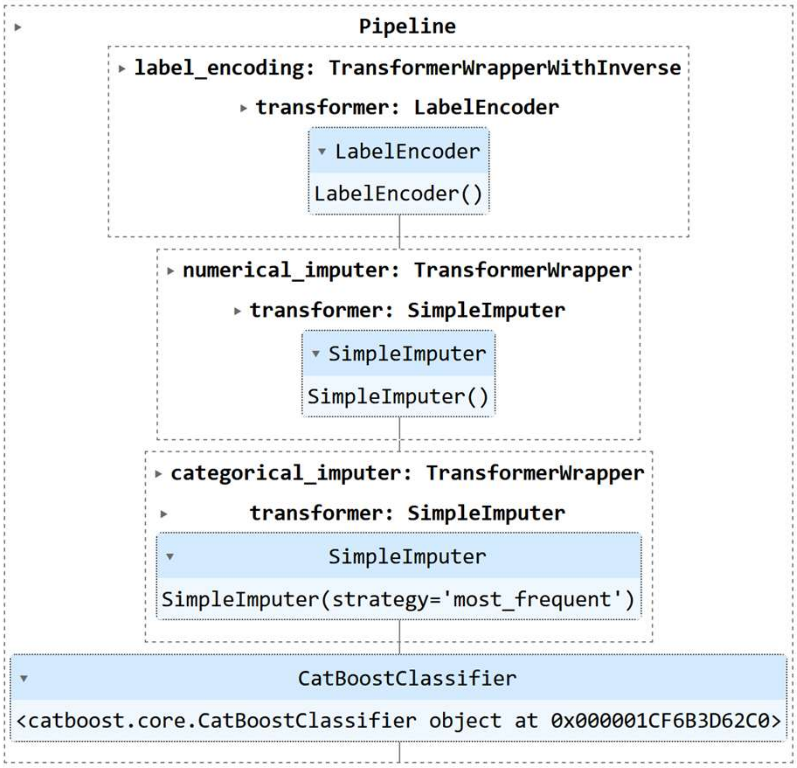


**Figure S1. PyCaret ML pipelines for impact prediction using eDNA.** A) MLP model; B) CatBoost model.

A) **MLP pipeline: classification accuracy 83.3%**

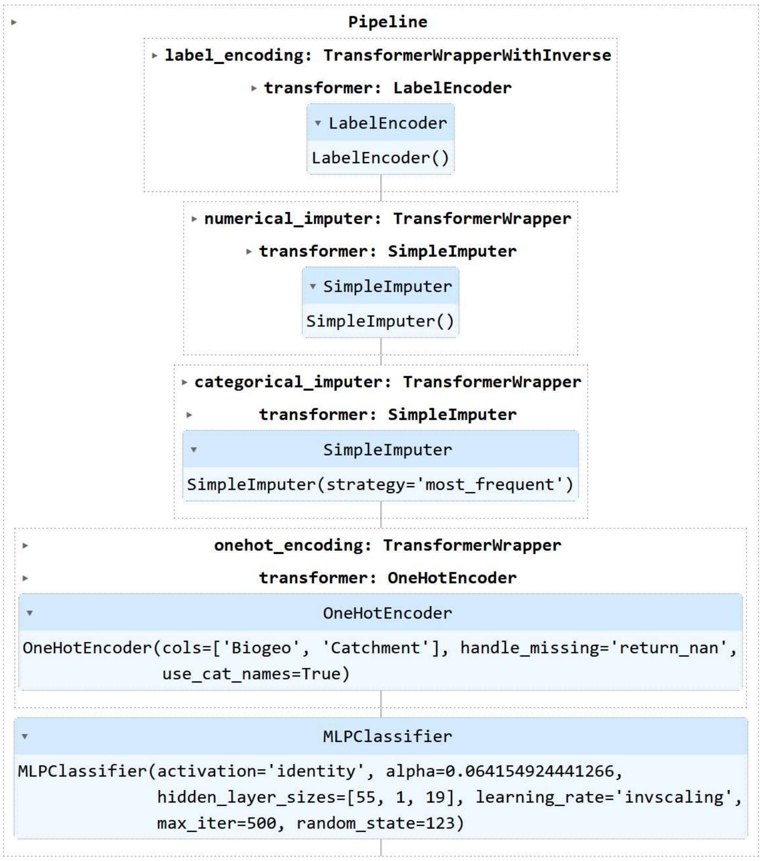


B) **CatBoost pipeline: classification accuracy 75.0%**

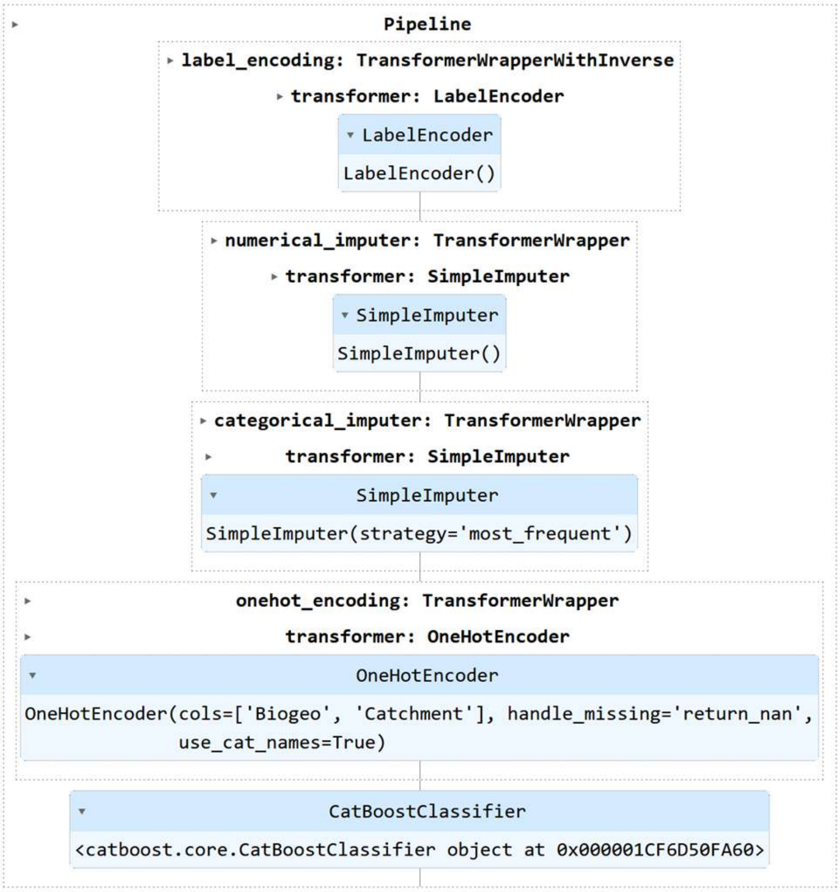


**Figure S2. PyCaret ML pipelines for impact prediction using combined eDNA and biogeographic data.** A) MLP model; B) CatBoost model.

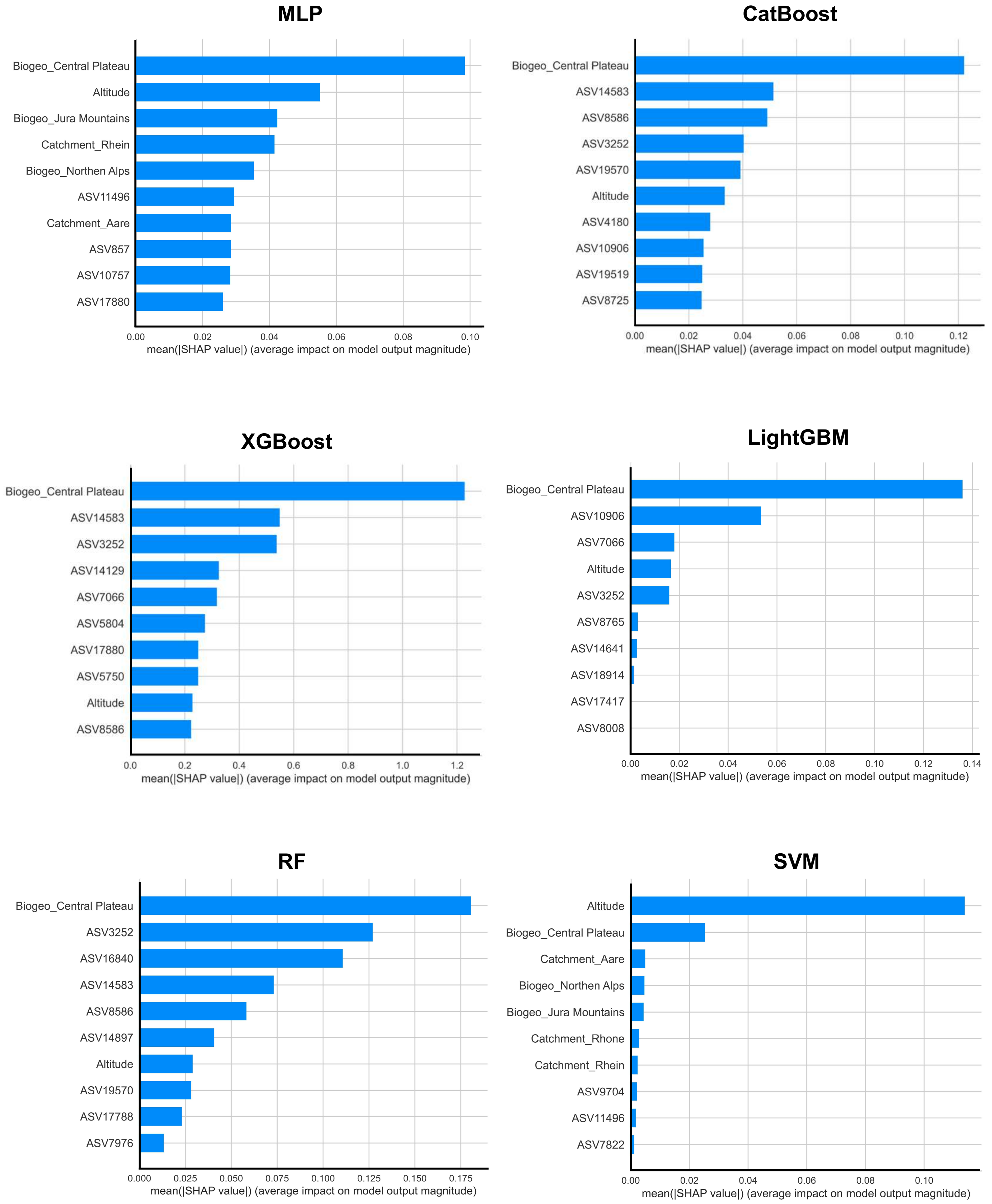


**Figure S3. SHAP bar plots showing top 10 features for six ML models using combined multimodal dataset for impact prediction.** Average impact of a feature on model prediction is summarized as mean absolute SHAP value.

**Table S2. Classification report for multimodal ML on test set**

| Model | Class | Precision | Recall | F1-Score | Support | Accuracy |
|---|---|---|---|---|---|---|
| MLP | Imp | 0.8333 | 0.8333 | 0.8333 | 24 | 0.8333 |
| | Ref | 0.8333 | 0.8333 | 0.8333 | 24 | |
| CatBoost | Imp | 0.7308 | 0.7917 | 0.7600 | 24 | 0.7500 |
| | Ref | 0.7727 | 0.7083 | 0.7391 | 24 | |
| XGBoost | Imp | 0.7391 | 0.7083 | 0.7234 | 24 | 0.7292 |
| | Ref | 0.7200 | 0.7500 | 0.7347 | 24 | |
| LightGBM | Imp | 0.7000 | 0.8750 | 0.7778 | 24 | 0.7500 |
| | Ref | 0.8333 | 0.6250 | 0.7143 | 24 | |
| RF | Imp | 0.7391 | 0.7083 | 0.7234 | 24 | 0.7292 |
| | Ref | 0.7200 | 0.7500 | 0.7347 | 24 | |
| SVM | Imp | 0.7500 | 0.5000 | 0.6000 | 24 | 0.6667 |
| | Ref | 0.6250 | 0.8333 | 0.7143 | 24 | |

Models were trained using the PyCaret AutoML package to predict impacted (Imp) or reference (Ref) status for biomonitoring sites using a combined dataset of eDNA and biogeographic data. Classification results for unseen data (test set) are presented.

**Table S3.** **Ecological significance of the top 10 ASV features in MLP for impact prediction using eDNA according to literature**

| ASV ID | Species Assigned | Feature Ranking | Impact on Reference | Ecological Significance in Literature | Reference |
|---|---|---|---|---|---|
| ASV11496 | *Baetis rhodani* | 1 | Negative | One of the most widespread mayfly species, with increased abundance in urban areas compared to natural environments and tolerant against heavy metal pollution. | 1,2,3 |
| ASV10757 | *Orthocladius telochaetus* | 2 | Positive | *Orthocladius telochaetus* is usually found in artic streams where temperatures > $2^0$C but <$4^0$C, indicating that it is sensitive to warming effects caused by anthropogenic activities. | 4,5 |
| ASV18087 | *Thienemanniella cf. vittata* | 3 | Positive | *Thienemanniella cf. vittata* was found in low relative abundance in nutrient enriched streams in contrast to other chironomids (*Eukiefferiella claripennis* and *Cricotopus bicinctus*) which had significant correlation with algal assemblages. | 6 |
| ASV17880 | *Simulium variegatum* | 4 | Positive | *Simulium vareigatum*, a type of black fly, is found to have reduced abundance in heavily polluted urban streams compared to natural environments. | 7 |
| ASV16981 | *Chironomidae sp.* | 5 | Positive | --- | --- |
| ASV13657 | *Orthocladiinae sp.* | 6 | Positive | --- | --- |
| ASV13720 | *Diamesa insignipes* | 7 | Negative | *Diamesa insignipes* is usually found in cold mountain streams is one of the most eurythermal (tolerant against a range of temperature) species within the genus *Diamesa,* suggesting some tolerance against warming effects caused by anthropogenic activities. | 8 |
| ASV15320 | *Orthocladiinae sp.* | 8 | Positive | --- | --- |
| ASV9710 | *Orthocladius frigidus* | 9 | Negative | Study suggests *Orthocladius frigidus* is tolerant to moderate levels of heavy metal (e.g., zinc) pollution in rivers. | 9 |
| ASV1493 | Unknown | 10 | Negative | --- | --- |

Top 10 ASV features and their association with the reference prediction were provided by SHAP analysis.

**References**


1. Bohus, A.; Gál, B.; Barta, B.; Szivák, I.; Karádi-Kovács, K.; Boda, P.; Padisák, J.; Schmera, D. Effects of urbanization-induced local alterations on the diversity and assemblage structure of macroinvertebrates in low-order streams. Hydrobiologia 2023, 850 (4), 881-899. DOI: 10.1007/s10750-022-05130-1.
2. G., R.; W., S. Heavy Metal Accumulation by Baetis rhodani and Macrobenthic Community Structure in Running Waters of the N' Harz Mountains (Lower Saxony/FRG) (Ephemeroptera: Baetidae). Entomologia Generalis 1991, 16 (1), 31-37. DOI: 10.1127/entom.gen/16/1991/31.
3. Vilenica, M.; Brigić, A.; Sartori, M.; Mihaljević, Z. Microhabitat selection and distribution of functional feeding groups of mayfly larvae (Ephemeroptera) in lotic karst habitats. Knowl. Manag. Aquat. Ecosyst. 2018, (419), 17.
4. Marziali, L.; Gozzini, M.; Rossaro, B.; Lencioni, V. Drift patterns of Chironomidae (Insecta, Diptera) in an Arctic stream (Svalbard Islands): an experimental approach. Studi Trentini di Scienze Naturali 2009, 84.
5. Stur, E.; Ekrem, T. The Chironomidae (Diptera) of Svalbard and Jan Mayen. Insects 2020, 11 (3). DOI: 10.3390/insects11030183.
6. Maasri, A.; Fayolle, S.; Gandouin, E.; Garnier, R.; Franquet, E. Epilithic chironomid larvae and water enrichment: is larval distribution explained by epilithon quantity or quality? Journal of the North American Benthological Society 2008, 27 (1), 38-51.
7. Ciadamidaro, S.; Mancini, L.; Rivosecchi, L. Black flies (Diptera, Simuliidae) as ecological indicators of stream ecosystem health in an urbanizing area (Rome, Italy). Ann Ist Super Sanita 2016, 52 (2), 269-276.
8. Rossaro, B. Factors that determine chironomidae species distribution in fresh waters. Bollettino di zoologia 1991, 58 (3), 281-286.
9. Ruse, L.; Ruse, L.; Herrmann, S.; Sublette, J. Chironomidae (Diptera) species distribution related to environmental characteristics of the metal-polluted Arkansas River, Colorado. Western North American Naturalist 2000, 34-56.

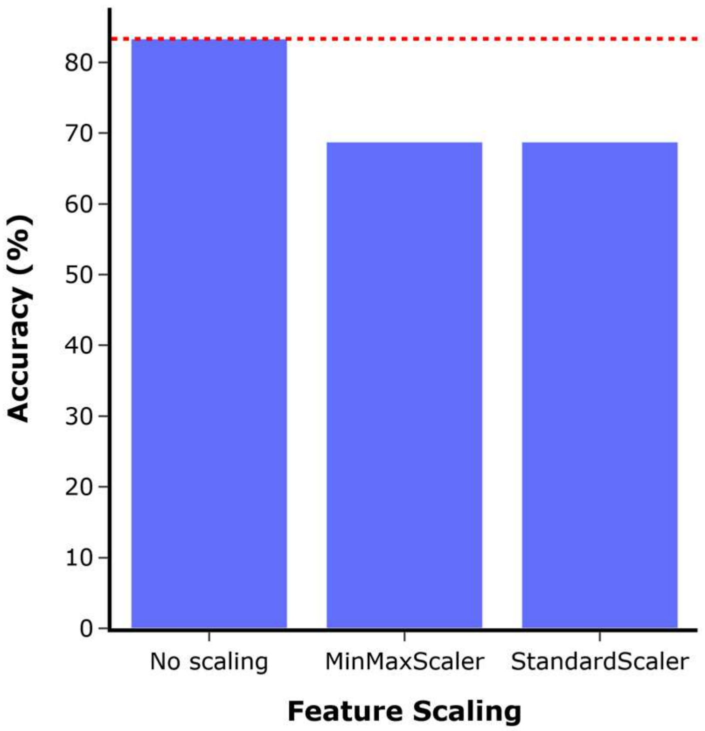


**Figure S4. Effect of altitude feature scaling on MLP performance for impact prediction using combined multimodal dataset.** The altitude feature was scaled using either the MinMaxScaler (values of 0 to 1) or StandardScaler which normalizes values using the Z-score. Both scaling methods are commonly applied for feature scaling in ML and are standard functions available in the Scikit-learn Python package. The best classification performance is indicated by red dashed line.
